# Proneural genes form a combinatorial code to diversify neocortical neural progenitor cells

**DOI:** 10.1101/2023.07.29.551096

**Authors:** Alexandra Moffat, Ana-Maria Oproescu, Satoshi Okawa, Sisu Han, Lakshmy Vasan, Hussein Ghazale, Daniel J Dennis, Dawn Zinyk, François Guillemot, Antonio del Sol, Carol Schuurmans

## Abstract

Neocortical neural progenitor cells (NPCs) are molecularly heterogeneous, yet the genes that confer distinct neuronal morphologies and connectivities during development are poorly understood. Here, we determined that a proneural gene combinatorial code diversifies cortical NPCs. By mining scRNA-seq data from murine embryonic and early postnatal cortices and generating trajectory inference models, we found that Neurog2 is predominant, and is transiently co-expressed with Ascl1 and/or Neurog1 during an apical-to-basal NPC transition state in NPCs with early pseudotime identities. To assess whether proneural gene pairs confer distinct properties, we first used Neurog2/Ascl1 reporter mice expressing unique reporters, revealing that NPCs have distinct cell division modes and cell cycle dynamics dependent on their proneural gene profile. To assess Neurog2/Neurog1 interactions, we used double knock-out mice and novel split-Cre transgenics crossed to a Rosa-diptheria-toxin-A line to delete double^+^ cells, showing Neurog1/Neurog2 are specifically required to generate early-born neurons and to maintain NPCs. Finally, in silico mutation of a cortical Neurog2-gene regulatory network and validation using Neurog1/Neurog2 mutant and ‘deleter’ mice, identified Bclllb and Nhlh2, expressed in early-born neurons, as dependent on Neurog1/Neurog2. Our data explains how proneural genes act combinatorically to diversify gene regulatory networks, thereby lineage restricting NPCs and creating cortical neuronal diversity.

## INTRODUCTION

The neocortex is a mammalian-specific brain region that is associated with higher level cognitive functioning and sensory processing. It is comprised of six layers of functionally-specialized, excitatory pyramidal neurons that display a range of diverse transcriptional signatures, connectivities and morphologies (Tasic et al. 2018). With the advent of single cell technologies, the molecular heterogeneity of the neural progenitor cells (NPCs) that build the neocortex has been revealed (Telley et al. 2016; Telley et al. 2019; Di Bella et al. 2021), yet the functional significance of this heterogeneity is not completely understood. The current dogma suggests that lineage-specifying transcription factors (TFs) induce neurogenesis and specify neuronal subtype identities, whereas signal-dependent TFs drive neuronal maturation (Alvarez-Dominguez and Melton 2022). Three proneural genes, Neurog1, Neurog2 and Ascl1, encode lineage-specifying basic-helix-loop-helix (bHLH) TFs that are expressed in cortical NPCs (Oproescu et al. 2021). As drivers of neurogenesis and determinants of cell identities, the proneural genes likely shape neuronal diversification (Oproescu et al. 2021). Neurog2 is the dominant neuronal determinant in the embryonic neocortex since it is both necessary and sufficient to promote the differentiation of glutamatergic neurons, and can largely compensate for the loss of Neurog1 or Ascl1 (and not vice versa) (Schuurmans et al. 2004; Britz et al. 2006; Li et al. 2012; Kovach et al. 2013; Dennis et al. 2017). The importance of Neurog2 as a central neuronal fate determinant is further supported by predictive transcriptomic studies (Telley et al. 2016). Nevertheless, Neurog1 is essential, as it is required to temper the pace of early neurogenesis by reducing the potent neurogenic activity of Neurog2 (Han et al. 2018). Furthermore, Ascl1 is also essential as it participates with Neurog2 to control Notch signaling in a subset of cortical niche cells, ensuring an even pattern of neurogenesis across the germinal zone (Han et al. 2021).

Neocortical glutamatergic neurons arise from dorsal telencephalic/pallial NPCs between embryonic day (E) 11 and 17 in mice (Takahashi et al. 1999; Caviness et al. 2009). The first-born cortical neurons arise from NPCs in the pallial margins, migrating tangentially to form a marginal zone (layer 1) populated by Cajal-Retzius (CR) neurons (Bielle et al. 2005). Subsequent rounds of neurogenesis lead to the sequential, inside-out generation of radially-migrating subplate (transient layer 7), deep-layer (layers 5, 6), and upper layer (layers 2-4) neurons (Villalba et al. 2021). Neurogenic NPCs are divided into apical and basal compartments (Vaid and Huttner 2022). The principal apical NPCs are apical radial glia (aRG), which reside in the ventricular zone (VZ) and extend processes to the ventricular and pial surfaces, with the basal extensions supporting glial-guided neuronal locomotion (Marín et al. 2010). Generally, aRG undergo direct neurogenesis during early corticogenesis, while later in development, aRG give rise to a basal pool of intermediate progenitor cells (IPCs) (Haubensak et al. 2004; Miyata et al. 2004; Noctor et al. 2004). IPCs lose their apical and basal contacts and move basally to form a subventricular zone (SVZ), where they divide 1-2 times before undergoing neurogenesis (Haubensak et al. 2004; Miyata et al. 2004; Noctor et al. 2004). Basal radial glia (bRG), which lose their apical and retain their basal processes, are expanded in species such as humans and other larger mammals that have gyrencephalic cortices (Hansen et al. 2010; Betizeau et al. 2013; LaMonica et al. 2013; Martinez-Martinez et al. 2016), yet only represent only ∼5% of the rodent basal NPC pool (Shitamukai et al. 2011; Martinez-Cerdeno et al. 2012).

A progressive fate restriction model was initially put forth to explain how the six layers of neocortical neurons are sequentially generated, positing that cortical NPCs gradually become fate-restricted to upper-layer neuronal fates through a series of temporal identity transitions (McConnell and Kaznowski 1991; McConnell 1995; Desai and McConnell 2000). Indeed, active transcriptional programs in cortical NPCs diversify over time, switching from an early phase of neurogenesis that is predominantly driven by intrinsic factors, to later phases of neurogenesis when NPCs become “exteroceptive”, or influenced by extrinsic environmental cues (Telley et al. 2016; Telley et al. 2019). This progressive restriction model also gained support from Cre-based lineage tracing experiments that followed the progeny of the earliest neurogenic NPCs; Sox9^+^ aRG (Kaplan et al. 2017) and Fezf2^+^ aRG (Guo et al. 2013), to reveal wide-spread laminar labelling. Conversely, other studies have supported the existence of fate-restricted cortical NPCs. For instance, Emx2^+^ (Garcia-Moreno et al. 2012) and Cux2^+^ (Franco et al. 2012) aRG lack neurogenic potential early in development, undergoing delayed neurogenesis during the window of upper-layer formation. However, these findings are controversial since other groups have found that Cux2^+^ aRG give rise to all cortical layers (Eckler et al. 2015; Gil-Sanz et al. 2015). Nevertheless, taken together, these studies support the idea that cortical NPCs are molecularly heterogeneous. Additional supporting evidence includes the demonstration that adult neural stem cells (NSCs) are derived from a small pool of embryonic aRG that are set aside in dormancy in the early embryo (Fuentealba et al. 2015; Furutachi et al. 2015).

Molecular heterogeneity of cortical NPCs is observed at the level of gene expression (i.e., high or low levels) and in the specific genes expressed. For instance, the amount of Sox9 expression dictates whether NPCs will self-renew or differentiate (Fabra-Beser et al. 2021). Furthermore, cortical NPCs that differentially express the proneural genes Neurog2 and Ascl1 have distinct transcriptomic and epigenomic signatures, and neurogenic versus gliogenic differentiation biases (Han et al. 2021). By stratifying cortical cells based on proneural gene expression using single-cell (sc) RNA-seq data (Di Bella et al. 2021), we found that Neurog2 is expressed throughout cortical lineage development, and co-expressed with Neurog1 and Ascl1 only in minor subsets of NPCs with early pseudotime identities that are transiting from an apical to basal cell fate. To decipher differences in cortical NPCs expressing different proneural gene combinations, we first used dual reporter mice to show that cortical NPCs expressing Neurog2 and/or Ascl1 differ in their modes of cell division and proliferation dynamics. We then used double knock-outs (DKO) and split-Cre ‘deleter’ mice to show that Neurog1/Neurog2 are required together to maintain cortical NPCs and for the differentiation of early-born cortical neurons. Finally, in silico mutation analyses revealed that Neurog1/Neurog2 use a unique gene regulatory network (GRN) to support the differentiation of early-born cortical neurons, which we validated with our deleter and DKO mice. Taken together, these data support the notion that proneural genes act in a combinatorial fashion to diversify neocortical neurons.

## RESULTS

### Neurog2 is co-expressed with Neurog1 and/or Ascl1 in small subpopulations of apical and basal NPCs

We previously mined transcriptomic data from human fetal tissue and cerebral organoids to demonstrate that NEUROG2 is the predominant proneural gene in human aRG, bRG and IPCs, while only a subset of these cells co-expressed ASCL1 (Han et al. 2021). Here, we asked whether murine cortical NPCs could similarly be distinguished by distinct patterns of proneural gene expression. For this purpose, we mined scRNA-seq data collected from mouse cortices between E10.5 and postnatal day (P) 4 (Di Bella et al. 2021). Using Seurat clustering, we created high dimensional UMAP plots, revealing the sequential appearance of ∼20 main cell clusters (Figure 1A,B). Cluster identities were assigned using cell type-specific markers for each cortical cell type as described (Di Bella et al. 2021) (Figure S1A), with main clusters corresponding to aRG, IPCs, layer (L) 1-6 neurons, interneurons, oligodendrocytes and astrocytes (Figure 1A,B).

**Figure 1.**
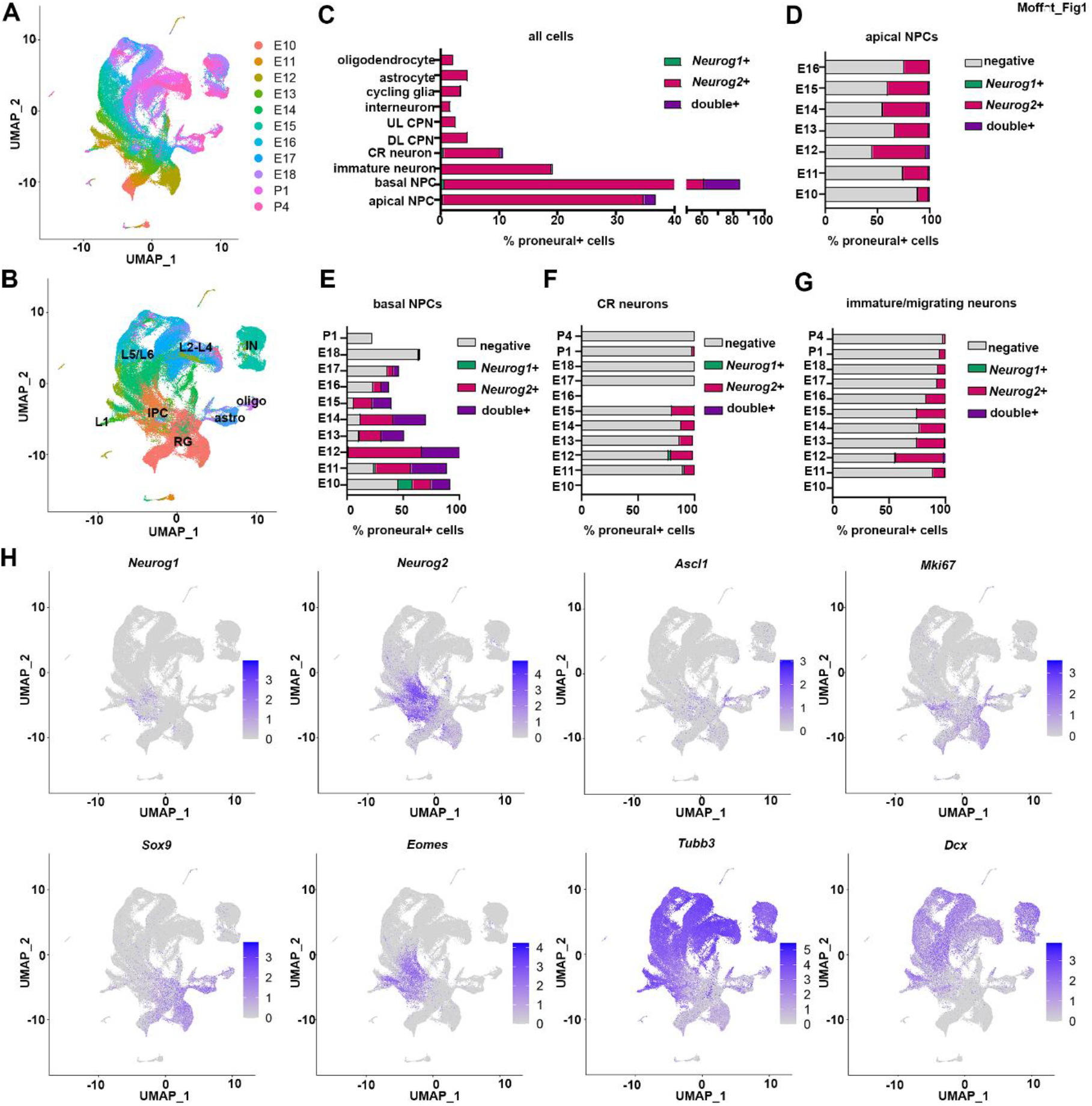
Mining scRNA-seq data to identify proneural gene-expressing cortical cells. (A,B) UMAP plot of scRNA-seq data from cortical tissue collected between E10 and P4 (GSE153162) (Di Bella et al. 2021), showing cell clusters associated with time points (A) and cell types (B). (C-G) Distribution of cell types expressing Neurog1 alone (green bars), Neurog2 alone (pink bars), co-expressing Neurog1 and Neurog2 (purple bars) or not expressing Neurog1 or Neurog2 (grey bars). Data is depicted as the total % of each cell population expressing the proneural genes (C), or separated into cell types, including apical NPCs (D), basal NPCs (E), Cajal-Retzius neurons (F) and immature and migrating neurons (G). (H) Feature plots depict the cortical populations expressing Neurog1 and Neurog2 and canonical markers of different cortical cell types. Astro, astrocyte; CR, Cajal-Retzius; DL, deep-layer; IN, interneuron; IPC, intermediate progenitor cell; L, layer; NPC, neural progenitor cell; oligo, oligodendrocyte; RG, radial glia; UL, upper layer. See also Figure S1 and S2.

Using these cluster assignments, we observed that IPCs most frequently expressed the proneural genes, beginning at E10 and persisting until P1 (Figure 1C,E). Within the IPC pool, 83.8% of cells expressed Neurog2, of which, 23.2% co-expressed Neurog1, and 9.0% co-expressed Ascl1 (Figure 1C,E; Figure S2A). IPCs that expressed Neurog1 (0.6%) or Ascl1 (1.0%) alone were infrequent (Figure 1C,E). The next cell population with the highest frequency of proneural gene expression was apical NPCs, of which 36.3% of were defined by Neurog2 expression, including 2.0% that co-expressed Neurog1 and 6.3% that co-expressed Ascl1 (Figure 1C,D; Figure S2A). Apical NPCs that only expressed Neurog1 (0.4%) or Ascl1 (2.8%) were once again in the minority (Figure 1C,D; Figure S2A). The bias towards Neurog2, Neurog1 and Ascl1 expression in basal NPCs was also evident in feature plots of the scRNA-seq data (Figure 1H; Figure S2C). Mapping Neurog1, Neurog2 and Ascl1 expression profiles onto the UMAPs revealed a striking similarity to feature plots showing Eomes expression, a canonical IPC marker (Figure 1H; Figure S2B). In contrast, there was less overlap between proneural gene feature plots compared to those depicting the expression of markers for dividing aRG (Mki67, Sox2, Sox9), immature neurons (Tubb3, Dcx), early-born (Reln, Bcll1b, Tbr1) or late-born (Satb2) neurons (Figure 1H).

In addition to being expressed in cortical NPCs, proneural gene transcripts were also detected in smaller subsets of differentiated cells. For instance, 10.5% of CR neurons expressed Neurog2, including 0.6% and 0.9% that co-expressed Neurog1 and Ascl1, respectively, whereas only a minor fraction of CR cells expressed either Neurog1 (0.7%) or Ascl1 (1.3%) alone (Figure 1C,F; Figure S2A). Additionally, 18.8% of migrating and immature neurons expressed Neurog2, including 0.3% and 0.4% that co-expressed Neurog1 and Ascl1, respectively (Figure 1C,G; Figure S2A). Neurog2 was also expressed in subsets of deep-layer (DL) and upper-layer (UL) cortical projection neurons (CPN) and interneurons, as well as in small proportions of cycling glia, oligodendrocytes and astrocytes in the late embryonic and early postnatal period (Figure S2C). Conversely, Ascl1 expression was more frequently detected in interneurons, cycling glia, oligodendrocytes and astrocytes, and in small numbers of DL and UL CPN beginning at P4 (Figure S2A). In summary, Neurog2 is the predominant proneural gene during murine corticogenesis and is expressed in most basal IPCs and in smaller numbers of apical RG, subsets of which co-express Neurog1 and Ascl1.

### Neurog1-Neurog2 lineage trajectory analyses

Neurog2 biases cortical NPCs towards SVZ-directed, non-surface divisions (Miyata et al. 2004), suggesting Neurog2 may be expressed during the transition phase from an apical to basal NPC. Previous trajectory inference analyses of NEUROG2/ASCL1 expressing cells in human cerebral organoids revealed a lineage bifurcation between neuronal and glial lineages (Han et al. 2021). Herein, we focused on the developmental trajectory of murine cortical NPCs expressing Neurog2 and/or Neurog1. We generated trajectory inference models from cortical cells that expressed one or both genes, using scRNA-sequencing data from E10.5 to P4 (Di Bella et al. 2021) (Figure 2A). Five cell states were identified, with state 1 having the ‘youngest’ pseudotime and state 5 the ‘oldest’ (Figure 2B,C). Mapping proneural gene expression onto these trajectories confirmed that cells rarely solely expressed Neurog1, except at the earliest pseudotimes in state 1 and state 2 (Figure 2D). In contrast, Neurog2^+^ and Neurog1/Neurog2 double^+^ cells were detected in all 5 pseudotime states, although double^+^ NPCs were enriched in earlier states (Figure 32D).

**Figure 2.**
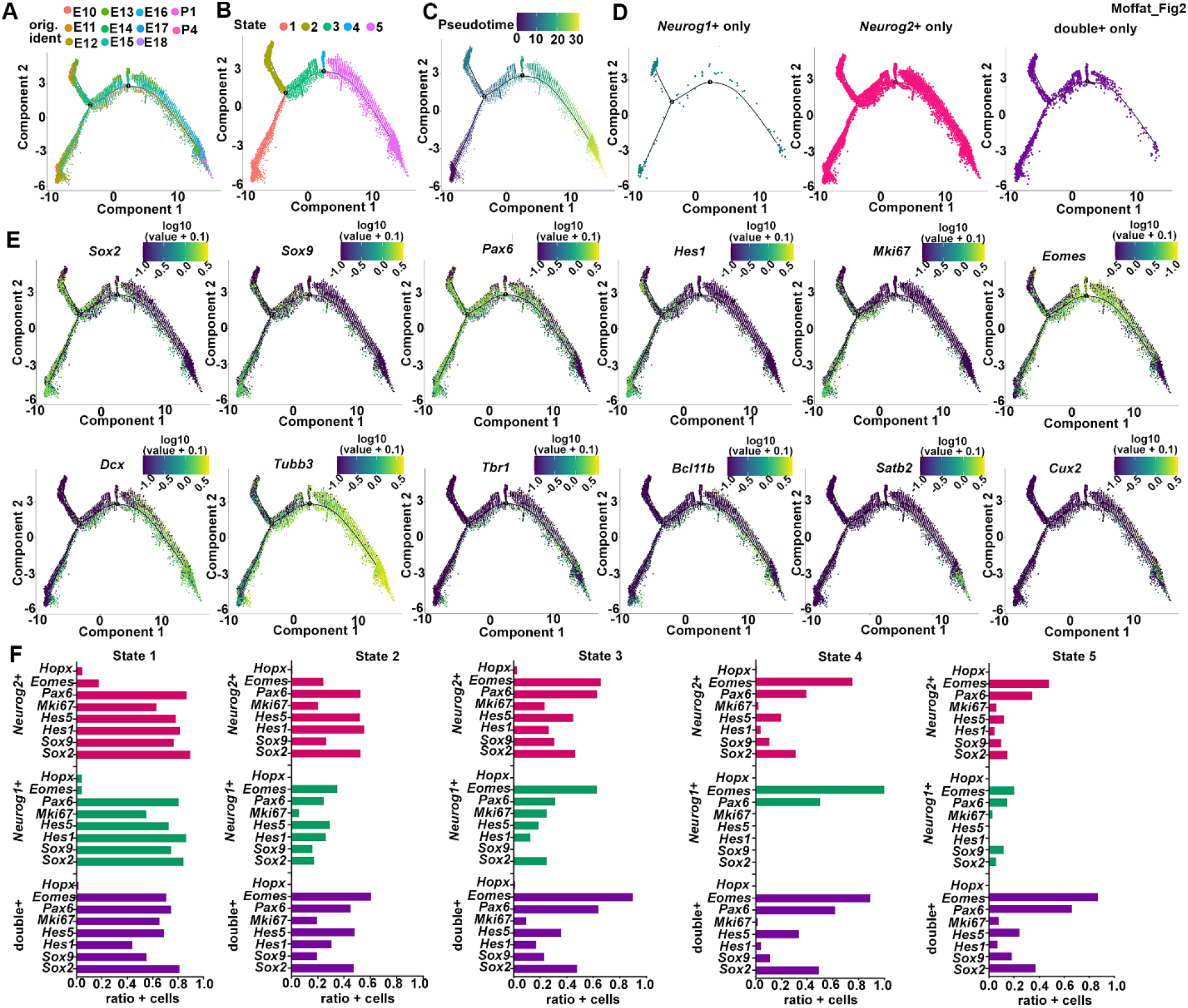
Pseudotime trajectory analysis of Neurog1 and Neurog2 expressing cortical cells. A-D) Monocle3 lineage trajectory analysis of Neurog1^+^, Neurog2^+^ and double^+^ cells collected between E10-P4 (A), showing annotation of states 1-5 (B), a pseudotime trajectory of these states (C), and the distribution of each NPC population in each state (D). (E,F) Mapping marker expression onto the pseudotime trajectory plots (E) and showing expression levels graphically (F). State 1 and 2 are predominated by apical NPC markers, state 3 and the minor state 4 populations are mainly IPCs, and state 5 are IPCs and newborn neurons. Neurog1^+^ (green bars), Neurog2^+^ (pink bars) and co-expressing (purple bars) cells predominate in earlier pseudotime states (F). See also Figure S3.

To identify cell populations within each state, we mapped the expression of known markers onto the pseudotime trajectories (Figure 2E). Neurog1 and Neurog2 expression was most abundant in pseudotime states 1 and 2 (Figure 2D, Figure S3A), which were enriched in the expression of the IPC marker, Eomes, along with aRG markers Sox2, Sox9, Pax6, and Hes1 (Figure 2E). Hopx, a bRG marker (Pollen et al. 2015; Johnson et al. 2018), was expressed at low levels in state 1 only (Figure 2E). At the earliest pseudotime in state 1, cells expressing Neurog1, Mki67, a proliferation marker, and aRG markers were elevated (Figure 2E,F). Conversely, in the earliest pseudotime of state 2, Neurog1^+^ cells were predominant, but Mki67 and aRG markers were detected in fewer cells, suggesting that state 1 and state 2 NPCs differ not only in pseuodotime but in their stem and proliferative potential (Figure 2E,F). Taken together, these data suggest that Neurog1/Neurog2 co-expressing cortical NPCs resemble aRG at early pseudotime states, and some of these cells express Eomes.

A lower proportion of Neurog1/Neurog2-expressing cells were detected in states 3 and 4 (Figure S3A), and of these, state 4 was a minor subpopulation (Figure 2B,E). States 3 and 4 both included a preponderance of cells expressing the IPC marker, Eomes, but there was a reduced frequency of aRG marker expression, and an increase in the ratio of cells expressing neuronal markers, such as Rbfox3 and Tubb3 (Figure S3B). Thus, even though NPCs that express Neurog1/Neurog2 have a similar transcriptional profile to Eomes^+^ IPCs (see UMAP plots, Figure 1H), most double^+^ and Neurog1^+^ NPCs have an earlier pseudotime identity compared to state 3 and 4 Eomes^+^ IPCs. Finally, Neurog2 was the only proneural gene detected at prominent levels in state 5, the latest pseudotime state (Figure S3A). State 5 was enriched in Eomes, Dcx, Tubb3, Tbr1, Bcllb, Satb2 and Cux2 expressing cells, with the latter two upper-layer markers detected in cells at the latest pseudotimes (Figure S3B). These data suggest that Eomes^+^ cells are stratified into an early cell identity characterized by Neurog1/Neurog2 and aRG marker co-expression, and a later state, in which the expression of proneural genes and other aRG markers declines and neuronal markers begin to be expressed.

### Neurog2 and Ascl1 single^+^ and double^+^ NPCs display distinct biases in cell division modes and unique cell cycle dynamics

To assess the functional significance of gene expression heterogeneity in cortical NPCs, several groups have used differential-labeling based on patterns and levels of gene expression, followed by assays for self-renewal, proliferation and/or differentiation (García-Moreno and Molnár 2015; Fabra-Beser et al. 2021; Han et al. 2021). Our working model was that double^+^ NPCs co-expressing two proneural genes have emergent properties that their single^+^ NPC counterparts would not. Using in vitro assays and in vivo lineage tracing, we previously showed that E12.5 Neurog2/Ascl1 single^+^ and double^+^ cortical NPCs have an increased propensity to exit the cell cycle compared to proneural-negative NPCs, and display distinct neuronal and oligodendroglial lineage biases (Han et al. 2021). To further distinguish Neurog2^+^, Ascl1^+^ and double^+^ NPCs, we first examined cell division modes using a pair cell assay (Dave et al. 2011). CD15^+^ Neurog2 negative (-) and positive (+) NPCs were sorted from E13.5 Ascl1^GFPKI/+^;Neurog2^FLAG-mCherryKI/+^ cortices and plated at clonal density (Figure 3A). At 24-hrs post-plating, we examined two cell clones by immunolabeling with β3-tubulin (TuJ1) to mark neurons and Ki67 to label dividing NPCs (Figure 1B,C). The proneural-negative NPC pool showed an ∼4-fold increase in symmetric proliferative divisions compared to all proneural^+^ NPCs (p<0.0001 in all pairwise comparisons; Figure 3D), mimicking the enhanced proliferative capacity of these cells characterized in a previous neurosphere assay (Han et al. 2021). In contrast, Neurog2^+^, Ascl1^+^ and double^+^ NPCs were biased towards symmetric differentiative divisions (∼4-fold increase compared to proneural-negative; p<0.0001 in all pairwise comparisons; Figure 3D), consistent with the ability of sustained Neurog2 or Ascl1 expression to promote neurogenesis (Imayoshi et al. 2010; Imayoshi et al. 2013). Ascl1^+^ and Neurog2^+^ NPCs were, however, distinguishable from each other, with Ascl1^+^ NPCs displaying a 3.4-fold increase in the proportion of asymmetric cell divisions compared to Neurog2^+^ NPCs (p=0.0081; Figure 3D). Thus, proneural gene-expressing cortical NPCs largely switch to symmetric neurogenic divisions, at least in vitro, regardless of whether they express Neurog2 or Ascl1.

**Figure 3.**
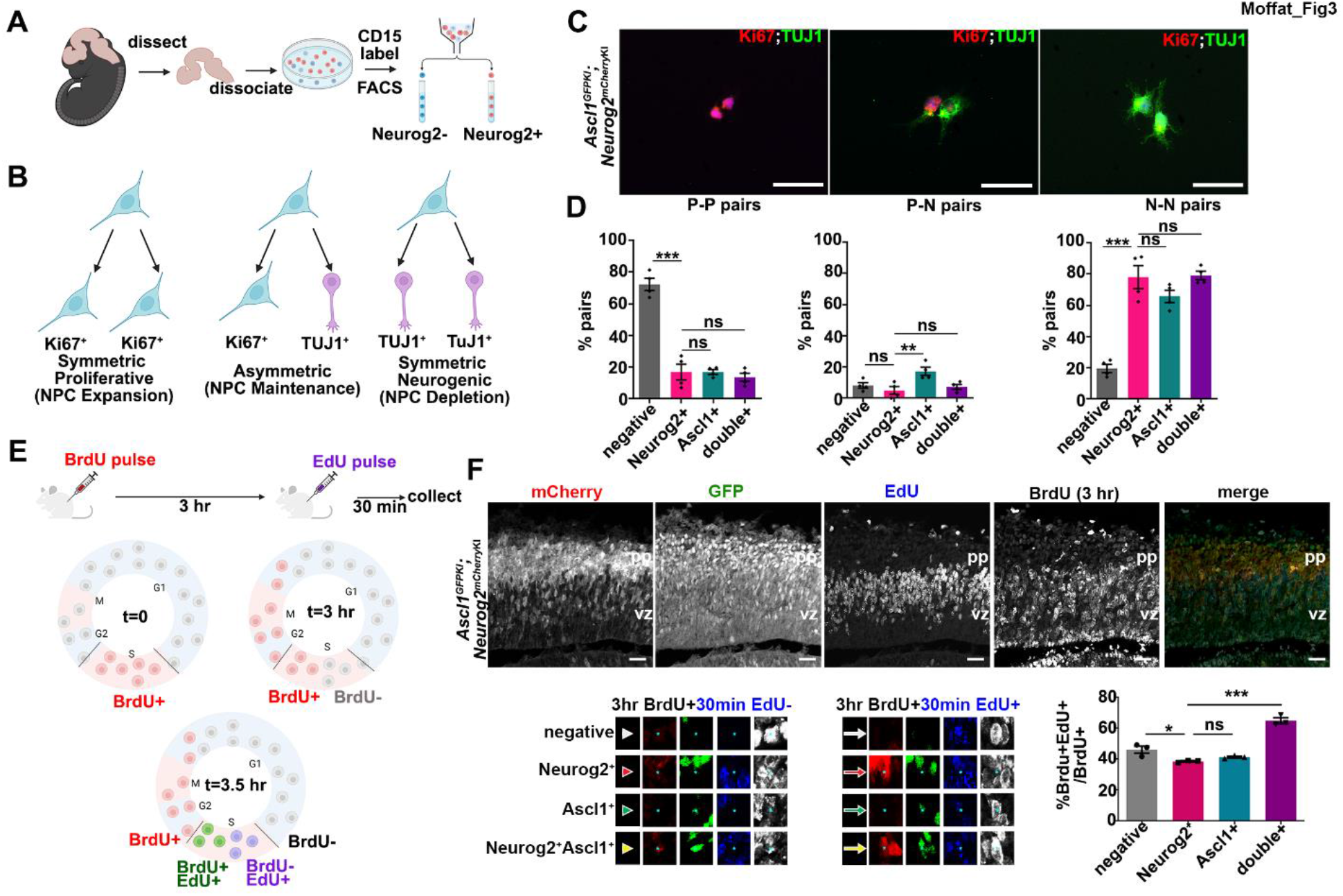
Neurog2/Ascl1 single^+^ and double^+^ cortical NPCs display differences in the mode and dynamics of cell division. (A,B) Schematic of pair-cell assay procedure (A) showing predicted cell division pattern outcomes (B). (C,D) Representative images of symmetric proliferative (two Ki67^+^ cells), asymmetric (one Ki67^+^/one β3-tubulin^+^), and symmetric differentiative (two β3-tubulin^+^ neurons) divisions. E12.5 FACS enriched NPCs (CD15^+^) from Neurog2^mCherryKI^;Ascl1^GFPKI^ cortices were separated into proneural negative, Neurog2^+^, Ascl1^+^ and double^+^ NPCs before plating (C). Quantitation of two cell clones showing symmetric proliferative (P-P pairs), asymmetric renewing (P-N pairs) and symmetric neurogenic (N-N pairs) divisions (N=4) (D). Scale bars = 25 μm. (E) Schematic of injection strategy for dual pulse labeling with BrdU and EdU. (F) Co-immunolabeling of mCherry, GFP, BrdU (3 hrs) and EdU (30 mins), and quantification of %BrdU^+^EdU^+^/BrdU^+^ cells (N=3, n=9). High magnification images of BrdU^+^EdU^-^ cells are shown below; white (negative), red (Neurog2^+^), green (Ascl1^+^) and yellow (double^+^) arrows indicate BrdU^+^/EdU^+^ cells. Scale bars in F indicate 100 μm. Data represent the mean ± s.e.m, p-values: ns - not significant, <0.05 *, <0.01 **, <0.001 ***, by one-way ANOVA with Tukey correction for multiple comparisons. pp, preplate; vz, ventricular zone.

Neural stem and progenitor cell populations differentiate asynchronously with cell cycle remodeling, especially G1 lengthening, observed in differentiating cells (Dalton 2015). To further distinguish the proliferation dynamics of cortical NPCs based on Neurog2/Ascl1 expression, we used a dual pulse labeling technique with nucleoside analogs to label all cells entering S-phase of the cell cycle in a 3 hr period, as a proxy measure of cell production rates. We first administered 5-bromo-9-deoxyuridine (BrdU) into E13.5 Neurog2^FLAG-mCherryKI/+^;Ascl1^GFPKI/+^ embryos, and after 3 hrs, a second thymidine analogue, 5-ethynyl-2′-deoxyuridine (EdU), was injected (Figure 3E). We focused on NPCs that were in S-phase for both pulses (i.e., %EdU^+^BrdU^+^/BrdU^+^) as a measure of entry into S-phase. The normal S-phase length in E13.5 cortical progenitors is 3.9 hrs (Takahashi et al. 1995) therefore, not all NPCs should pick up both S-phase labels. Neurog2^+^ NPCs had a 1.2-fold lower EdU^+^BrdU^+^/BrdU^+^ ratio than proneural negative NPCs (p=0.0301; Figure 3F), consistent with the idea that fewer of these cells re-enter the cell cycle. In contrast, entry of double^+^ NPCs into S-phase was elevated 1.7-fold compared to Neurog2^+^ NPCs (p<0.0001). Taken together, these studies indicate that Neurog2^+^ NPCs are not only molecularly distinct from Ascl1^+^ and Neurog2/Ascl1 double^+^ NPCs (Han et al. 2021), but also display distinct biases in their dynamics of cell division.

### Neurog1 and Neurog2 are required to maintain aRG morphology and to prevent the precocious depletion of cortical NPCs

The heterogeneous nature of the Neurog2/Ascl1 expressing NPC pool prompted us to ask whether Neurog2^+^ NPCs could be further stratified by their co-expression with a different proneural gene, Neurog1. We did not have access to Neurog1 and Neurog2 transgenic mice with different fluorescent reporters to isolate and track single^+^ and double^+^ NPCs. Instead, we relied on KO mice to examine the genetic requirement for Neurog1 and Neurog2 in cortical NPCs. Previous DKO studies revealed that Neurog1 and Neurog2 are required to specify the glutamatergic neuronal phenotype of early-born, deep-layer cortical neurons (Fode et al. 2000; Schuurmans et al. 2004). We first asked whether the loss of Neurog1/Neurog2 impacts aRG, the primary NPCs. From double heterozygous intercrosses collected at E15.5, DKOs were distinguishable by prominent neural tube curvature and histologically by cortical plate thinning (Figure S4A,B). BLBP/Fabp7 and RC2/Nes immunolabeling of aRG revealed largely parallel radial processes that extended from the ventricular to pial surface in E15.5 wild-type cortices (Figure S4C). In contrast, in E15.5 DKOs, aRG were less organized and whorls of prematurely truncated fibers were observed near the ventricle (Figure S4C). By E18.5, the DKO phenotype was more severe, with the basal processes of aRG no longer arranged in parallel bundles and most fibers not reaching the pial surface (Figure 4A). In E18.5 Neurog2 KO cortices, most aRG contacted the pial surface, although some fibers prematurely terminated in a subpial layer, whileNeurog1 KO aRG appeared similar to wild-type (Figure 4A). Neurog1/Neurog2 are thus together required to retain the basal connection of aRG (Figure 4B), the loss of which is normally associated with aRG to IPC transitions (Borrell and Gotz 2014).

**Figure 4.**
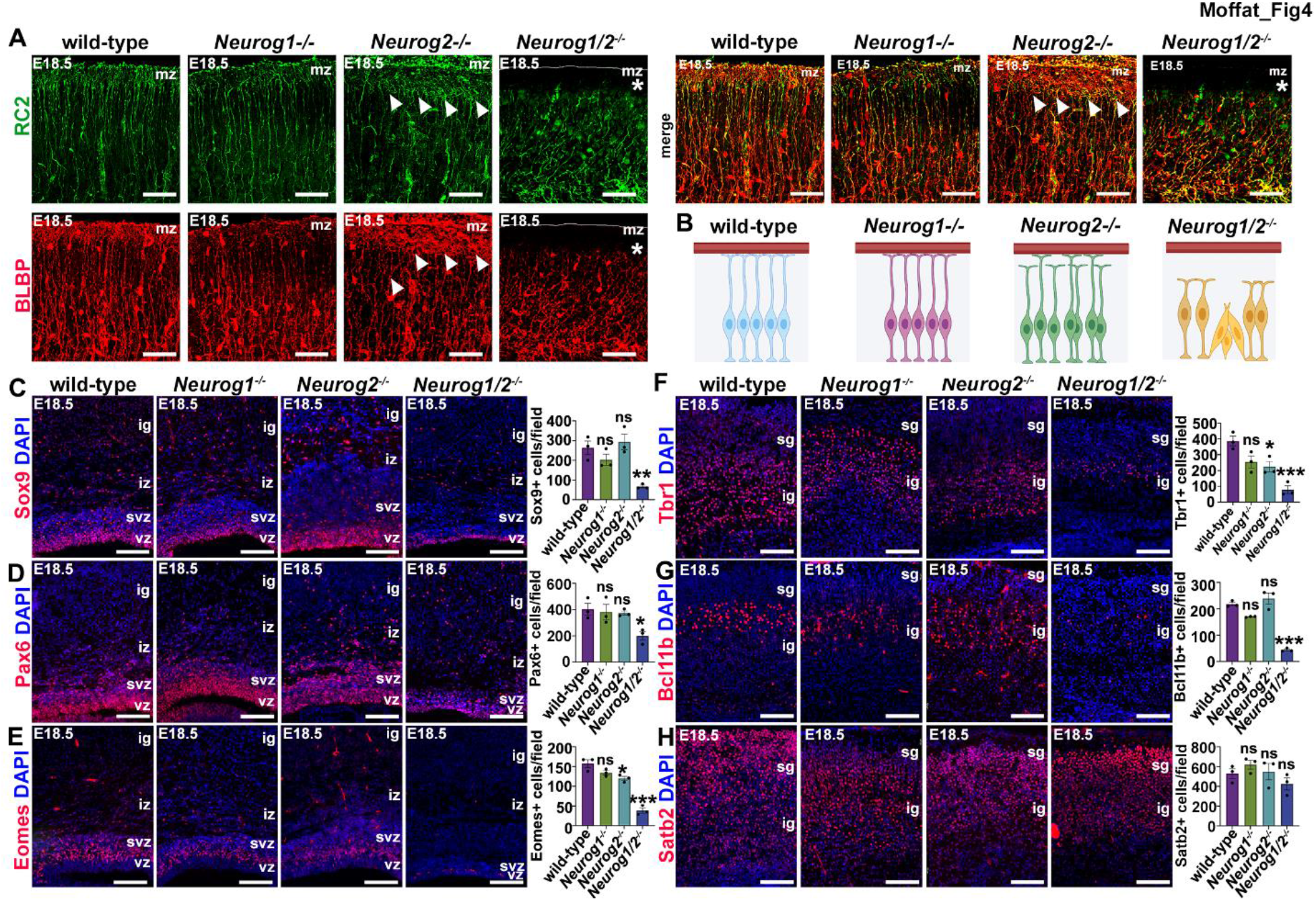
Radial glial defects, precocious depletion of NPCs, and reduced production of deep-layer neurons in Neurog1;Neurog2 DKO cortices. (A,B) RC2 and BLBP immunostaining of the aRG scaffold in E18.5 cortices revealed Neurog1;Neurog2 DKO aRG basal processes are disorganized and do not reach the pial surface (*), and more minor aberrant terminations of the basal endfeet in Neurog2 KOs (arrowheads) (A). Schematic representation of defects in radial glial guides in Neurog1/Neurog2 KO and DKO cortices (B). (C-H) Analysis of Sox9^+^ NPCs (C), Pax6^+^ NPCs (D), Eomes^+^ NPCs (E), Tbr1^+^ neurons (F), Bcl11b^+^ neurons (G) and Satb2^+^ neurons (H) in E18.5 wild-type, Neurog1^-/-^, Neurog2^-/-^ and Neurog1^-/-^;Neurog2^-/-^ cortices. Graphs show quantification of markers^+^ cells/field (N=3, n=9 for all genotypes). Scale bars are 25 μm in A and 100 μm in C-H. Blue is DAPI counterstain. NPC markers were only counted in the germinal zone. Data represent the mean ± s.e.m, p-values: ns - not significant, <0.05 *, <0.01 **, <0.001 ***, by one-way ANOVA with Tukey correction for multiple comparisons. cp, cortical plate; gz, germinal zone; mz, marginal zone; ig, infragranular layers; iz, intermediate zone; sg, subgranular layers; svz, subventricular zone; vz, ventricular zone. See also Figure S4.

To determine whether there was an acceleration of the aRG to IPC transition in DKOs, we quantified apical and basal NPCs in single KOs and DKOs, using Sox9 and Pax6 to mark aRG and Eomes to label IPCs. At E18.5, after neurogenesis is complete, there was a sharp reduction in Sox9^+^ aRG (3.9-fold, p=0.0071; Figure 4C), Pax6^+^ aRG (2.0-fold, p=0.030; Figure 4D) and Eomes^+^ IPCs (4.1-fold; p<0.0001; Figure 4E) only in DKOs, indicative of a precocious depletion of both apical and basal NPCs. To assess the impact of this premature NPC loss, we asked whether neurogenesis was altered. Both Tbr1^+^ (4.9-fold; p=0.0006; Figure 4F) and Bcl11b^+^ (5.0-fold; p<0.0001; Figure 4G) deep-layer neurons were reduced in DKO cortices, while upper-layer Satb2^+^ neurons were unaffected (Figure 4H). Thus, in the absence of Neurog1 and Neurog2, the cortical NPC pool is precociously depleted, however, only deep-layer neurogenesis is impacted.

### Generating split-Cre transgenics to lineage trace Neurog1/Neurog2 double^+^ NPCs

The Neurog1/Neurog2 DKO phenotype is an amalgamation of the loss-of-function of these genes in NPCs that individually express and co-express both genes. To specifically characterize the roles of Neurog1/Neurog2 within double^+^ NPCs, we devised a novel split-Cre lineage tracing system in which C-Cre was knocked into the Neurog1 locus (this study) and N-Cre was knocked into the Neurog2 locus (Han et al. 2021) (Figure 5A). Both Cre termini were fused to the coiled-coil domain of the eukaryotic transcriptional activator GCN4, which promotes intermolecular associations (Hirrlinger et al. 2009b). In Neurog1^C-CreKI^;Neurog2^N-CreKI^ transgenics (hereafter split-Cre mice), when a cell co-expresses the two genes, N- and C-Cre termini dimerize to re-constitute an active Cre. To validate our split-Cre system, we first confirmed the expression of the appropriate Cre termini in proneural^+^ NPCs by performing RNAScope in situ hybridization on E12.5 cortices heterozygous for Neurog1^C-CreKI^ or Neurog2^N-CreKI^ alleles, revealing a high coincidence level of Neurog1 and C-Cre, and Neurog2 and N-Cre transcripts, matching the endogenous expression of Neurog1 and Neurog2 (Figure 5B,C). To identify Neurog1/Neurog2 double^+^ NPC progeny, we performed lineage tracing in P150 split-Cre;Rosa-zsGreen brains. zsGreen^+^ cells were predominately located in forebrain structures, including in all layers of the neocortex, hippocampal CA1 and CA3 fields and dentate gyrus, the olfactory bulb, thalamus and amygdala (Figure S5A). Within the P150 neocortex, 38.1±4.8% of zsGreen^+^ cells expressed Satb2, while only 4.36±0.74% of zsGreen cells co-labeled with the glial marker Sox9 (N=3, Figure S5B,C). This finding is consistent with the understanding that Neurog1/Neurog2-expressing NPCs largely giving rise to neuronal lineages.

**Figure 5.**
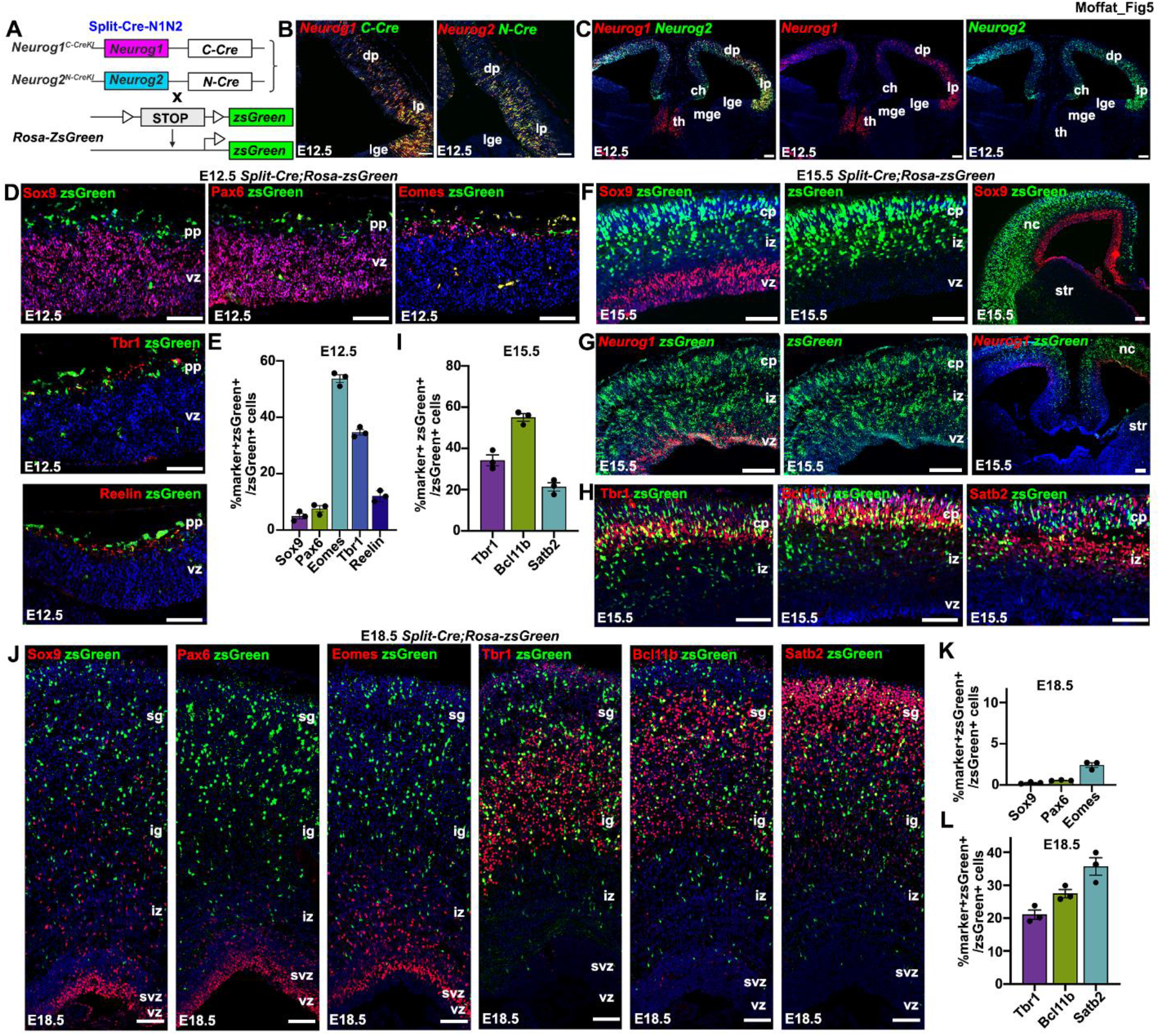
Split-Cre strategy to lineage trace Neurog1/Neurog2 co-expressing NPCs. (A) Neurog1^C-CreKI^;Neurog2^N-CreKI^ (split-Cre) strategy. (B,C) RNAscope in situ hybridization of Neurog1/C-Cre and Neurog2/N-Cre (B) and Neurog1/Neurog2 (C). (D,E) Co-labeling of zsGreen with Sox9, Pax6, Eomes, Tbr1 and Reelin in E12.5 split-Cre;Rosa-zsGreen cortices. (D). Quantification of zsGreen co-expression at E12.5 (N=3, n=9) (E). (F-I) Lineage tracing in E15.5 split-Cre;zsGreen cortices, showing co-labeling of zsGreen with Sox9 in low and high magnification images (F). Co-localization of zsGreen and Neurog1 mRNA in low magnification and higher magnification images (G). Co-labeling of zsGreen with Tbr1, Bcl11b and Satb2 in E15.5 split-Cre;zsGreen cortices (H). Quantification of of zsGreen co-expression at E15.5 (N=3, n=9) (I). (J-L) Co-labeling of zsGreen with Sox9, Pax6, Eomes, Tbr1, Bcl11b, and Satb2 in E18.5 split-Cre;zsGreen cortices (J). Quantification of zsGreen-NPC (K) and -neuronal (L) marker co-expression. (N=3, n=9 for all). (A-L) Blue is DAPI counterstain. Scale bars are 100 μm. Data represent the mean ± s.e.m, ch, cortical hem; dp, dorsal pallium; lge, lateral ganglionic eminence; lp, lateral pallium; mge, medial ganglionic eminence; nc, neocortex; pp, preplate; str, striatum; svz, subventricular zone; th, thalamus; vz, ventricular zone. See also Figure S5.

To assess the development of Neurog1/Neurog2 co-expressing cortical lineages, we examined split-Cre;Rosa-zsGreen transgenics throughout embryogenesis. At E12.5, most zsGreen^+^ cells co-labeled with Eomes, an IPC marker (53.69±1.38%), followed by Tbr1^+^ deep-layer neurons (34.67±1.00%) and Reelin (Reln)^+^ CR neurons (12.06±1.11%) (N=3, Figure 5D,E). In contrast, very few E12.5 Sox9^+^ (4.92±0.97%) and Pax6^+^ (7.56±1.11%) aRG expressed zsGreen (N=3, Figure 5D,E). Similarly, at E15.5, next to no zsGreen expression was detected in Sox9^+^ aRG (Figure 5F). Nevertheless, we confirmed that the Rosa-zsGreen locus was recombined in the VZ since zsGreen transcripts were present (Figure 5G). Thus, there is a time lag associated with a delay in the translation, maturation and/or accumulation of the fluorescent zsGreen protein. Consequentially, we may have under-estimated the true proportions of Neurog1/Neurog2 double^+^ NPCs. Nevertheless, we were able to use zsGreen protein for lineage tracing at E15.5 to show that double^+^ cells co-expressed markers for deep-layer Tbr1^+^ (28.19±1.98%) and Bcl11b^+^ (30.40±1.60%) neurons, and to a lesser extent, upper-layer Satb2^+^ neurons (15.78±0.69%) (N=3, Figure 5H,I). Finally, at E18.5, after neurogenesis is complete and gliogenesis has begun, very few zsGreen^+^ cells co-localized with Sox9 (0.08±0.04%), Pax6 (0.54±0.03%), or Eomes (2.40±0.28%) (N=3, Figure 5J,K). Ιnstead, double^+^ cells co-expressed Tbr1 (21.05±1.42%), Bcl11b (27.44±1.26%), and Satb2 (35.68±2.66) (N=3, Figure 5J,L). Thus, despite only being required for deep-layer neurogenesis, Neurog1/Neurog2 double^+^ NPCs give rise to both deep- and upper-layer neurons.

### Double^+^ NPCs are required to maintain the NPC pool and for deep-layer neurogenesis

To functionally test the requirement for double^+^ NPCs during cortical development, we crossed split-Cre mice with a Rosa-diphtheria toxin subunit A (DTA) transgenic line to selectively kill double^+^ NPCs (Figure 6A). Specifically, we used RC::L-DTA transgenic mice, which carries a floxed, inverted DTA gene in the Rosa locus that is only expressed upon Cre-mediated inversion, thus reducing the likelihood of leaky expression (Plummer et al. 2017). In triple transgenic split-Cre;RC::L-DTA mice, hereafter referred to as split-Cre;DTA mice, Cre should be fully re-constituted only in Neurog1/Neurog2 double^+^ NPCs. In double^+^ NPCS, the Rosa locus will be recombined to allow DTA expression, inducing cell death (Figure 6B). Since the RC::L-DTA allele contains an EGFP reporter under the ubiquitous control of the CAG promoter, GFP expression could be detected in all cortical cells of embryos carrying this allele (Figure 6C). In split-Cre;DTA ‘deleter’ cortices at E12.5, TUNEL^+^ apoptotic cells were enriched compared to littermate controls, especially in the preplate layer where postmitotic neurons reside (Figure 6C).

**Figure 6.**
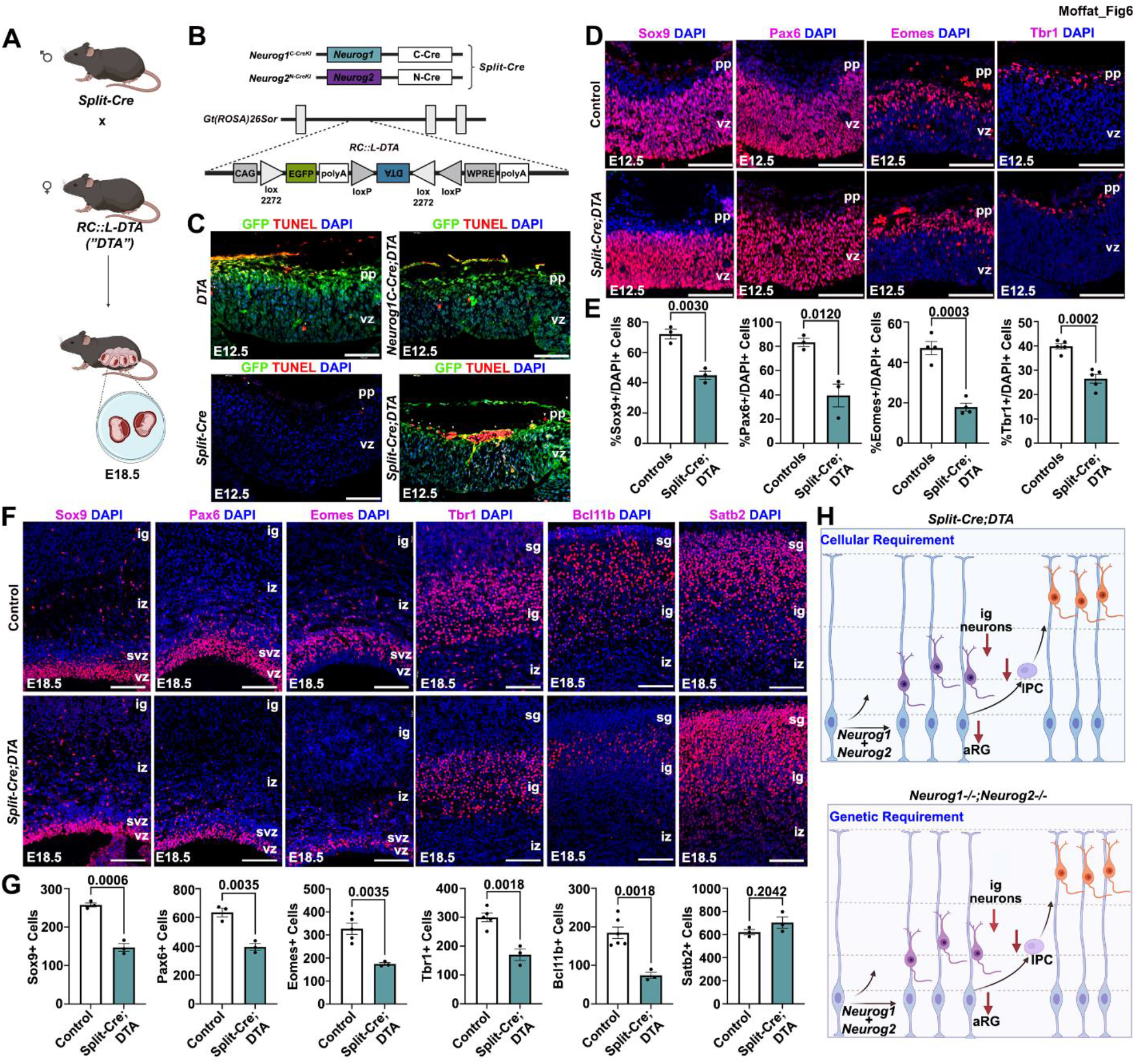
Double^+^ NPCs prevent precocious depletion and generate deep-layer neurons. (A,B) Crossing RC::L-DTA with split-Cre mice (split-Cre;DTA) to delete double^+^ cells (A). In split-Cre;DTA cells, Cre-mediated recombination flips the DTA transgene into its correct orientation (B). (C) E12.5 TUNEL^+^ apoptotic cells (red) accumulate in the preplate of split-Cre;DTA mice. (C-F) (D,E). E12.5 control and split-Cre;DTA cortices labelled with Sox9, Pax6, Eomes and Tbr1 (D). Quantification of marker^+^ cells/field: Sox9 (N=3, n=9 for both), Pax6 (N=3, n=9 for both), Eomes (N=4, n=12 for both) and Tbr1(N=5, n=15 for both) (E). (F,G) Sox9, Pax6, Eomes, Tbr1, Bcl11b and Satb2 immunostaining of E18.5 control and split-Cre;DTA transgenics (F). Quantification of marker^+^ cells/field: Sox9, Pax6, Satb2 (N=3, n=9 for both), Eomes, Tbr1 (N=5, n=15 for controls, N=3, n=9 for split-Cre;DTA), Bcl11b (N=6, n=18 controls, N=3, n=9 split-Cre;DTA)(G). (H) Cellular and genetic requirement for double^+^ NPCs. Blue is DAPI counterstain. Scale bars are 100 μm. Data represent the mean ± s.e.m, p-values: ns - not significant, <0.05 *, <0.01 **, <0.001 ***, by an unpaired Students t-test. ig, infragranular; iz, intermediate zone; sg, subgranular; svz, subventricular zone; vz, ventricular zone.

We next examined ‘deleter’ mice to determine how the loss of Neurog1/Neurog2 double^+^ NPCs impacted the overall NPC pool and cortical neurogenesis. In split-Cre;DTA ‘deleter’ cortices at E12.5, there was a reduction in the number of aRG expressing Sox9 (1.6-fold decrease; p=0.0030) and Pax6 (2.1-fold decrease; p=0.0120), and in IPCs expressing Eomes (2.6-fold decrease; p=0.0003) (Figure 6D,E). Further, we observed significantly fewer cells expressing the early neuron marker, Tbr1 (1.5-fold decrease; p=0.0002) (Figure 6D,E). In E18.5 triple transgenics, we observed a reduction in cells expressing all NPC markers: Sox9 (1.8-fold decrease; p=0.0006), Pax6 (1.6-fold decrease; p=0.0035) and Eomes (1.9-fold decrease; p=0.0035) (Figure 6F,G). Additionally, we found a significant reduction in Tbr1^+^ (1.8-fold decrease; p=0.0018) and Bcl11b^+^ (2.5-fold decrease; p= 0.0018) cells, markers of early-born L6 and L5 neurons, respectively (Figure 6F,G). In contrast, Satb2^+^ late-born, upper-layer neurons were detected in normal numbers in E18.5 triple transgenic cortices (p= 0.2042) (Figure 6F,G). Thus, the loss of cortical NPCs that co-express Neurog1 and Neurog2 affects deep-layer and not upper-layer neurogenesis. These data solidify the notion that Neurog1/Neurog2 are required in a distinct set of cortical NPCs that are destined to become deep-layer neurons (Figure 6H).

### In silico perturbation of Neurog1 and Neurog2 identifies co-regulated target genes

To identify target genes that may be co-regulated by Neurog1 and Neurog2, we examined the impact of in silico mutations on a cortical-specific, Neurog2-centered gene regulatory network (GRN) previously generated using RNA-seq and ATAC-seq data from E12.5 cortical NPCs that express Neurog2, and Neurog2 ChIP-seq data (Han et al. 2021) (Figure 7A). While the in silico ‘KO’ of Neurog1 was not predicted to disturb the Neurog2^+^ GRN, several genes were predicted to be downregulated after in silico ‘KO’ of Neurog2, as previously reported (Han et al. 2021) (Figure 7A). Moreover, in dual in silico ‘KO’ of Neurog1 and Neurog2, three new putative target genes were predicted to be downregulated; Neurod2, Bcl11b and Nhlh2 (Figure 8A). Notably, Neurod2 was previously shown to require Neurog1 and Neurog2 for its expression using a lower resolution Affymetrix microarray approach (Gohlke et al. 2008).

**Figure 7.**
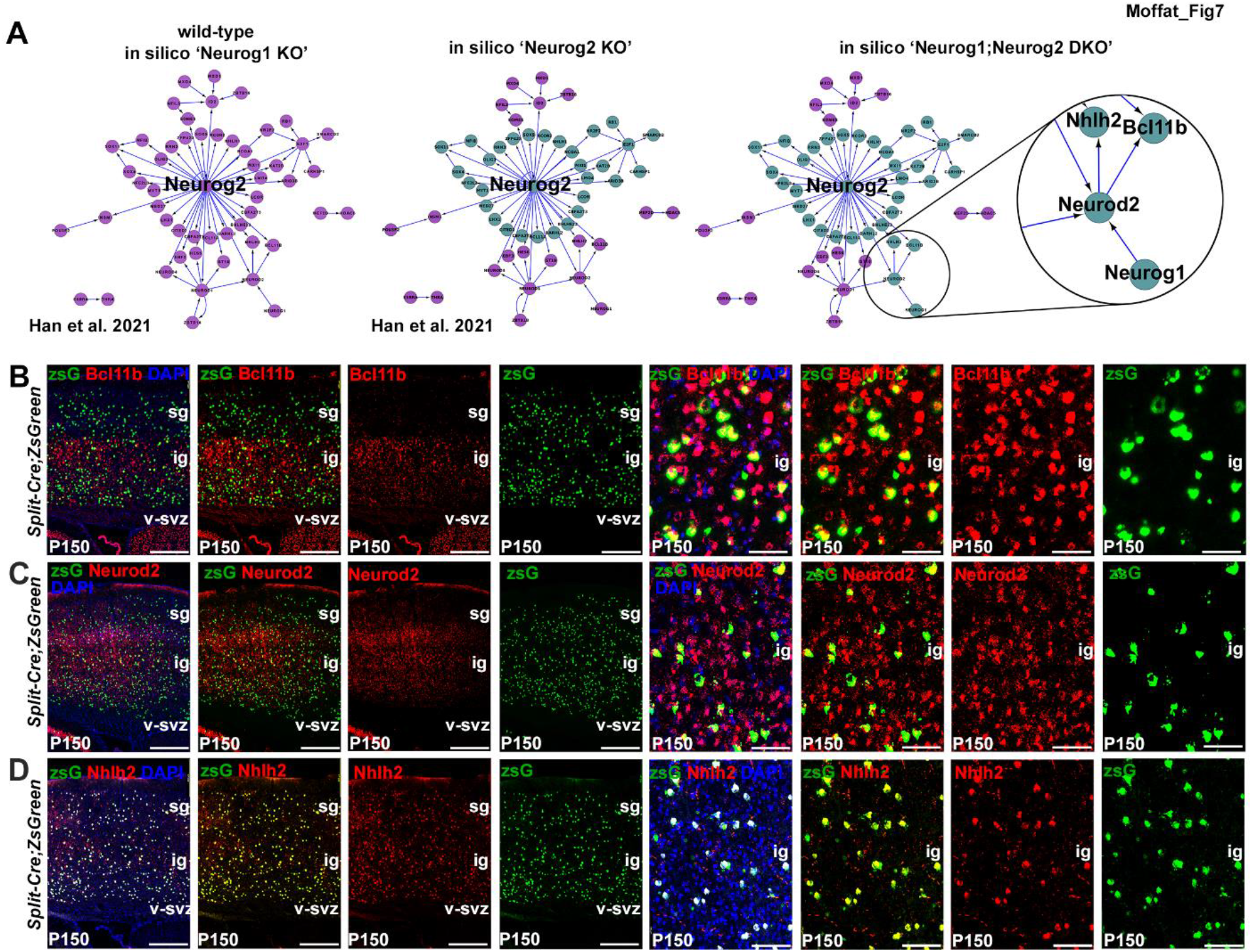
Identifying target genes co-regulated by Neurog1 and Neurog2 using gene regulatory network (GRN) and in silico mutation analyses. (A) GRN associated with E12.5 CD15^+^ Neurog2^+^ cortical NPCs, showing predicted target genes (purple nodes), from Han et al. 2021. Predicted changes in gene expression (teal nodes) caused by *in silico* ‘Neurog1 KO’ (no change, this study), Neurog2 (teal nodes, as generated in (Han et al. 2021)) and ‘Neurog1;Neurog2 DKO’, predicting four new target genes (teal nodes). (B-D) Co-expression of zsGreen with Bcl11b (B), Neurod2 (C) and Nhlh2 (D) in P150 split-Cre;Rosa-zsGreen cortices, with higher magnification images to the right. Blue is DAPI counterstain. Scale bars in B-D are 400μm on the left and 100μm on the right. ig, infragranular; iz, intermediate zone; pp, preplate. sg, supragranular; v-svz, ventricular subventricular zone. See also Figure S6.

Consistent with the idea that Neurod2, Bcl11b and Nhlh2 were indeed expressed in the Neurog1/Neurog2-dependent lineage, markers for all three genes co-localized with zsGreen^+^ cells in P150 split-Cre;Rosa-zsGreen reporter mice (Figure 7B-D; Figure S6A). This finding supplements our aforementioned observation that Bcl11b expression was reduced in E18.5 Neurog1/Neurog2 DKO cortices (Figure 4G) and E18.5 split-cre:Rosa-DTA deleter cortices (Figure 6F,G). Furthermore, we acquired a single surviving P150 split-Cre;DTA animal, which was overall smaller in size and lacked olfactory bulbs (Figure S6B), a feature of Neurog1/Neurog2 DKO embryos (Shaker et al. 2012). Notably, Bcl11b^+^ and Nhlh2^+^ cells appeared reduced in this P150 split-Cre;DTA cortex, whereas Neurod2^+^ and Satb2^+^ cells appeared unaffected (Figure S6C-E). Thus, while Neurod2, Bcl11b and Nhlh2 are expressed in cortical lineages derived from Neurog1/Neurog2 co-expressing NPCs, only Bcl11b and Nhlh2 appear to depend on these two proneural genes, at least partially, for their expression.

## DISCUSSION

To build an operational brain and to support brain plasticity, NPCs undergo adaptive changes in their proliferation dynamics and differentiation potential depending on their location (*e.g.,* lateral versus medial neocortex), embryonic-stage (*e.g.,* early versus late) and host species (*e.g.,* rodents vs. humans). While transcriptomic studies have revealed that NPCs are transcriptionally heterogeneous at any given stage (Telley et al. 2016; Telley et al. 2019; Di Bella et al. 2021), only certain genes drive cell differentiation and lineage diversification. Proneural genes specify neuronal phenotypes across animal taxa, making them ideal candidates to diversify neocortical neuronal lineages (Baker and Brown 2018). One of the big unknowns in the field of neocortical development has been why three proneural genes, namely Neurog1, Neurog2, and Ascl1, are expressed in embryonic cortical NPCs. Herein, we asked whether NPCs that express two proneural genes have emergent properties that are not observed in NPCs expressing a single proneural gene.

By mining scRNA-seq data collected from E10-P4 cortices, we found an enrichment of Neurog2, Neurog1 and Ascl1 transcripts in a cell cluster expressing Eomes, an IPC marker. In contrast, we found that fewer cells expressing canonical aRG markers such as Sox2, Sox9, Pax6 and Hes1, also co-expressed proneural genes. Trajectory inference modeling demonstrated that in comparison to Neurog2 single positive NPCs, NPCs co-expressing Neurog2 and either Ascl1 or Neurog1 had earlier pseudotime identities amongst the proneural^+^ NPCs. An overlap between Eomes and Pax6 expression has previously been reported in the basal half of the VZ (Englund et al. 2005), where Neurog2 expression is also enriched (Miyata et al. 2004; Britz et al. 2006). Our data thus supports a model in which proneural genes are expressed in transient Eomes/Pax6 co-expressing aRG that are transitioning to IPCs. After this transition, proneural gene and aRG marker expression shuts off, Eomes expression persists, and neuronal markers are upregulated. These data are consistent with a recent study performed in human iPSC-derived neuronal cultures that detected ASCL1 expression in a transient NPC state just prior to neuronal differentiation (Păun et al. 2023).

We identified differences in cell cycle dynamics and cell division modes between Neurog2/Ascl1 single^+^ and double^+^ NPCs, which supports the idea that cortical NPCs differ depending on which proneural gene(s) they express. These data are consistent with our previous demonstration that Neurog2/Ascl1 co-expression defines a distinct NPC pool that sustains a regular pattern of neurogenesis and prevents cortical folding by functioning as Notch-ligand expressing niche cells (Han et al. 2021). Cell cycle remodeling is known to drive asynchronous differentiation patterns of stem and progenitor cells within a tissue (Dalton 2015). The ability of Ascl1 to induce the expression of genes that promote cell cycle progression (Raposo et al. 2015) and Neurog2 to suppress pro-proliferative genes (Lacomme et al. 2012) exemplifies how proneural genes are excellent candidates to differentially remodel the cell cycle. Neurog2 activity is also restricted to G_1_ phase when differentiation programs are activated (Miyata et al. 2004; Britz et al. 2006). Tight regulation of proneural gene function is therefore required to prevent their inappropriate activity during the cell cycle. Several mechanisms regulate the differentiation potentials of Ascl1 and Neurog1/2 and may influence their cell cycle restricted activities, including phosphorylation (Ali et al. 2011; Hindley et al. 2012; Li et al. 2012; Li et al. 2014; Quan et al. 2016; Azzarelli et al. 2022), mRNA translation (Yang et al. 2014; Amadei et al. 2015; Zahr et al. 2018), protein degradation (Urban et al. 2016) and interactions with chromatin modulators (Seo et al. 2005; Lin et al. 2017; Păun et al. 2023).

We also investigated whether the co-expression of Neurog1 and Neurog2 confers specific properties onto cortical NPCs. Split-Cre lineage tracing suggested that Neurog1/Neurog2 double^+^ NPCs give rise to subsets of neurons in all cortical layers, similar to Neurog2/Ascl1 split-Cre lineage tracing (Han et al. 2021). However, to our surprise, Neurog1/Neurog2 are only required for the differentiation of a subset of deep-layer neurons and not upper-layer neurons. Notably, upper-layer neurons derived from Eomes^+^ and Eomes^-^ NPCs have distinct electrophysiological properties (Tyler et al. 2015), highlighting the importance of transcription factors in defining mature neuronal phenotypes. In the future, a similar approach could be used to determine whether neurons derived from Neurog2/Neurog1 double^+^ NPCs differ from those that are not derived from these NPCs, or instead derived from Neurog2/Ascl1 double^+^ NPCs.

Neurog2 and Ascl1 share some common functions including the induction of neurogenesis, specification of neuronal phenotypes, promotion of the apical to basal NPC transition (Miyata et al. 2004; Britz et al. 2006), control of neuronal migration (Hand et al. 2005; Ge et al. 2006; Heng et al. 2008; Pacary et al. 2011; Pacary et al. 2013), and the orientation of cortical NPC divisions (Pacary et al. 2013; Garcez et al. 2015). There are differences, however, since Neurog2 specifies a glutamatergic neuronal identity and Ascl1 specifies a GABAergic fate in the embryonic telencephalon (Oproescu et al. 2021). Furthermore, when Neurog2 and Ascl1 physically interact (Gradwohl et al. 1996; Henke et al. 2009; Han et al. 2021), they cross-repress one another, preventing the transactivation of their unique gene targets, and thus acting as competing lineage determinants (Han et al. 2021). Similarly, physical interactions between competing lineage determinants to prevent the transactivation of downstream targets has also been demonstrated in other tissues, as exemplified by GATA1 and PU.1 involved in priming hematopoietic lineages (Huang et al. 2007; Enver et al. 2009). Neurog1 can also heterodimerize with Neurog2, but it does not change the ability of Neurog2 to promote a glutamatergic neuronal fate, instead, Neurog1 modifies the pace of early cortical neurogenesis by reducing the potent neurogenic activity of Neurog2 (Han et al. 2018).

To achieve their distinct outcomes, Neurog2 and Ascl1 activate different downstream genes (Masserdotti et al. 2015), establish distinct epigenetic landscapes (Raposo et al. 2015; Aydin et al. 2019; Han et al. 2021), and use distinct GRNs (Han et al. 2021). Operationally, bHLH TFs heterodimerize. The combined expression of two proneural genes presents an opportunity for the formation of new heterodimers with unique functions, which is illustrated by Neurog2. Alone, Neurog2 is highly transcriptionally active and can efficiently induce neurogenesis, however, when combined with Neurog1 or Ascl1, it is less active (Han et al. 2018; Han et al. 2021). Unique heterodimers may bind different binding sites due to the preferred E-boxes specific to each proneural TF (Neurog2: CAKMTG, where K=G/T and M=A/C; Ascl1: CAGSTG, where S=G/C) (Wapinski et al. 2013; Raposo et al. 2015; Aydin et al. 2019). For example, the genes Dll1 and Dll3 are co-regulated by Neurog2 and Ascl1 in different ways.

While Dll1 has unique Neurog2 and Ascl1 binding sites in its regulatory region (Castro et al. 2006), Dll3 also has E-boxes that are specifically regulated by Neurog2-Ascl1 heterodimers (Henke et al. 2009). Neurog2 specifies a glutamatergic fate by initiating the expression of a battery of neuronal differentiation genes including direct downstream targets, Neurod1 and Eomes (Mattar et al. 2004; Schuurmans et al. 2004; Gohlke et al. 2008; Mattar et al. 2008; Ochiai et al. 2009). Here, we identified Bcl11b, Neurod2 and Nhlh2 as putative co-regulated genes of Neurog1 and Neurog2 and validated the dependence of Bcl11b and Nhlh2 on Neurog1/Neurog2 expression. Whether Neurog1 and Neurog2 bind as heterodimers to E-boxes in the upstream regulatory regions of these genes, or recognize distinct E-boxes, remains to be determined.

In summary, by stratifying cortical NPCs based on their combined expression of Neurog2-Ascl1 or Neurog2-Neurog1 expression, we have identified a primary mechanism for the differential control of cell cycle dynamics and neuronal fate specification during cortical development. Furthermore, we have identified a new population of lineage-restricted cortical NPCs (Neurog1/Neurog2 double^+^) that specifically give rise to early-born, deep-layer neurons, supporting the notion that there are unique cortical NPC pools with specific fates. Future analyses of the functional properties of these uniquely lineage-traced neurons will further our understanding of how cortical neuronal diversity arises.

## MATERIALS AND METHODS

### Animal sources and maintenance

All animal procedures were performed with the approval of the SRI Animal Care Committee (AUP 21-769) and followed the Canadian Council on Animal Care guidelines. Animals were housed in the Sunnybrook Research Institute (SRI) Comparative Research Unit (Toronto, ON, Canada) with ad libitum access to food and water. For expression analyses, wild-type CD1 (Charles River) females and males were interbred to generate embryos of the appropriate stage. The generation of Neurog1^GFPKI^ (Ma et al. 1998) and Neurog2^GFPKI^ (Britz et al. 2006) null mutant mice and Neurog2^Flag-mCherryKI/+^ reporter mice were previously described and maintained on a CD1 background (Han et al. 2021). The generation of Neurog2^N-CreKI^ mice were previously reported (Han et al. 2021) and the generation of Neurog1^C-CreKI^ transgenic mice is described below. The combined split-cre lines were crossed with Rosa-zsGreen (Jackson Lab: 007906) or RC::L-DTA (Jackson Lab: 026944) transgenics. Timed pregnancies were staged using the day of the vaginal plug as E0.5. Genotyping primers and PCR conditions are in the Supplemental Data section.

### Generation of transgenic mice

Neurog1^C-CreKI^ transgenic mice were generated by Cyagen using homologous recombination in embryonic stem cells and were produced on a BL6/J background (Jackson Labs). Neurog1^C-CreKI^ mice were created by replacing the Neurog1 coding domain with the C terminus of Cre (amino acids 60-343), also linked to a GCN4 coiled-coil domain as a flexible linker (Hirrlinger et al. 2009a; Beckervordersandforth et al. 2010). VectorBuilder (https://en.vectorbuilder.com) created the targeting vectors using 109-FUV-hGFAP-NCre and 106-FUV-P2-CCre sequences, as described (Hirrlinger et al. 2009a; Beckervordersandforth et al. 2010).

### Tissue processing

Pregnant females were euthanized by cervical dislocation following isoflurane anesthesia and embryos were dissected at embryonic stages. Embryos were removed from the uterine sac and brains were dissected with surgical forceps and blades in chilled 1X diethylpyrocarbonate (depc)-treated phosphate-buffered saline (PBS) over ice. Tail samples from each embryo were set aside for later genotyping. Brains were labeled and fixed overnight in 4% paraformaldehyde (PFA) diluted in 1X depc-PBS at 4°C. Brains were then rinsed 3×10 min in 1X depc-PBS and kept overnight in 20% sucrose diluted in 1X depc-PBS at 4°C. Embryonic brains were embedded in OCT on dry ice, stored at −80°C, and then cryosectioned at 10-micron thickness onto Superfrost™ Plus Slides (Fisherbrand™). All tools and processing materials were sterilized for RNAse-free working conditions, and samples were all wrapped in tin foil throughout processing to preserve transgenic reporter fluorescence. Adult mice were transcardially perfused with ice-cold 0.9% saline and 4% PFA in 1X depc-PBS. Brains were harvested and post-fixed in 4% PFA overnight at 4°C. As above, brains were rinsed 3×10 min in 1X depc-PBS and immersed in 20% sucrose diluted in 1X depc-PBS overnight at 4°C. Brains were embedded in OCT on dry ice and cryosectioned at 30-micron thickness onto Superfrost™ Plus Slides (Fisherbrand™). Slides were stored at −80°C until processing.

### RNAscope in situ hybridization

RNAScope^®^ technology (ACD Bio) was used for high-resolution (single transcript-level) detection of mRNA expression according to the manufacturer’s instructions in the RNAScope^®^ Multiplex Fluorescent Kit v2 User Manual (ACD #323110). Reagents were diethylpyrocarbonate (depc)-treated as required and RNAse-free working conditions were ensured. The following RNAScope^®^ probes were used: catalog versions of Mm-Neurog1 (Cat #414211), Mm-Neurog2 (Cat #417291), custom versions of N-Cre (#1062701-C3) and C-Cre (#1058921-C2) described in (Han et al. 2021), Mm-Nhlh2 (Ct #527811-C2), and a custom zsGreen1 probe (#895651-C3). Opal^TM^ 520 (Akoya #FP1487001KT; 1:1500), Opal^TM^ 570 (Akoya #FP1488001KT; 1:1500) and Opal^TM^ 690 (Akyoya #FP1497001KT; 1:1500) were used to stain channel 488 (green), 568 (red), or 647 (far-red). Slides were stored in the dark at 4°C until imaging.

### Immunostaining

Cryosections were rinsed 5 times for 5 minutes in 0.1% Triton X-100 in 1X PBS (1X PBST), incubated for 1 hour at room temperature in blocking solution (10% horse serum in 1X PBST) and then incubated overnight at 4°C with primary antibodies. Post-natal slides were rinsed 5 times for 5 minutes in 0.3% Triton X-100 in 1X PBS (0.3% PBST), incubated for 2 hours at room temperature in blocking solution (10% horse serum in 0.3% PBST) and then incubated overnight at 4°C with primary antibodies. For Bcl11b, a citrate antigen retrieval step was performed in a steamer using citrate buffer (pH=7.0) at a temperature of 97°C for 20 minutes. Following antigen retrieval, sections were washed 5 times for 5 minutes in 0.1% PBST and post-natal slides in 0.3% PBST. The following primary antibodies diluted in blocking solution were used: rabbit-anti-Sox9 (Millipore #AB5535, 1:300), rabbit-anti-Eomes (Abcam #ab23345, 1:300), rabbit-anti-Pax6 (BioLegend #901301, 1:300), rabbit-anti-Tbr1 (Abcam #ab31940, 1:300), mouseIgG1-anti-Satb2 (Abcam #ab51502, 1:300), rat-anti-Ctip2 (Abcam #ab18465, 1:500), mouseIgG1-anti-Bcl11b (Novus Biologicals #NB100-2600, 1:500), rabbitIgG-anti-Neurod2 (Abcam #ab104430, 1:500), mouseIgG1-anti-NeuN (Millipore Sigma, #MAB377, 1:500), rabbitIgG-anti-Neurog2 (Invitrogen #PA5-785, 1:500). Sections were then rinsed 5x for 5 minutes in 1X PBST and incubated for 1.5 hours at room temperature with secondary antibodies. Secondary antibodies were all diluted 1:500 in blocking solution and conjugated to different fluorophores, including Alexa Fluor™ 568 (red: rabbit IgG #A10042; mouse IgG1 #A21124), Alexa Fluor™ 555 (red: mouse IgG2a #A21137) or Alexa Fluor™ 647 (far-red: rabbit IgG, Invitrogen #A31573). Sections were then counterstained with 4’,6-diamidino-2-phenylindole (DAPI) (D1306, 1:10,000, Invitrogen) diluted in 1X PBST (1:10,000). Coverslips were mounted onto slides using Aqua-Poly/Mount (Polysciences, #18606) and left to dry overnight at 4°C. Slides were stored in the dark at 4°C until imaging.

### Histology, BrdU and EdU

For brain histology, cortices were dissected out at E15.5 and fixed in Bouin’s fixative overnight at 4°C before being rinsed and stored in 70% ethanol. Brains were then dehydrated, embedded in paraffin, sectioned at 7μm and stained with cresyl violet. To label S-phase cells, timed-pregnant females were injected intraperitoneally with 100 μg/g of BrdU or EdU as indicated. Brains were fixed and processed as described for cryosectioning. Prior to BrdU immunostaining, sections were treated with 2N HCl at 37°C for 20 minutes, followed by a PBST wash. Sections were then blocked in 10% horse serum in 0.3% PBST for 1 hr at room temperature before incubating in primary antibodies, including rat anti-BrdU (1/500, Serotec #OBT0030S). EdU injected tissue samples were stained by Click-iT® Plus EdU Imaging Kits (Molecular Probes, cat. C10640) using the manufacturer’s protocol.

### Radial glial immunostaining

Cortices were dissected out and fixed overnight in 2% PFA in PBS for 6 hrs at 4°C. Brains were washed 3 times in 1X PBS and then blocked in 3% low-melting point agarose prior to sectioning at 100 mm on a Leica VT1000S vibratome. Floating sections were permeabilized with PBST, and blocked for 1 hr in 10% heat-inactivated fetal bovine serum (FBS) in PBST. Sections were then incubated with primary antibodies overnight at 4°C, including rabbit anti-Blbp (1:500, Abcam #ab32423) and mouse anti-RC2 (nestin, 1/500, Developmental Studies Hybridoma Bank, AB_531887). Sections were then rinsed 5 times for 5 minutes in 1X PBST, incubated for 1.5 hrs at room temperature with secondary antibodies, and washed again 5 times in 1X PBST. Sections were then mounted onto slides using aquapolymount.

### Click-iT Plus TUNEL™ Assay

10μm cryosections were washed 5 times for 5 minutes in 0.1% PBST. Slides were fixed with 4% paraformaldehyde (PFA) for 15 minutes, incubated at 37°C then washed with 0.1% PBST for 5 minutes. Proteinase-K, provided by the Click-iT Plus TUNEL™ Assay Kit (Invitrogen #C10619) was used to permeabilize the tissue. Slides were washed for 5 minutes with 0.1% PBST, fixed with 4% PFA for 5 minutes at 37°C, and washed with 0.1% PBS-T and deionized water. A TdT reaction was performed followed by the Click-iT Plus™ reaction containing Alexa Fluor™ 647 picolyl azide dye, as indicated in the manufacturer’s guidelines. DNA was stained using Hoescht 33342™ solution. Sections were imaged with a Leica DMI8 fluorescent microscope and Hoescht 33342™ and Alexa Fluor™ 647 stains were detected with excitations of 350nm and 650nm, respectively.

### Pair cell assay

We performed a modified pair cell assay as described in (Dave et al. 2011). Briefly, brains from E12.5 Neurog2^Flag-mcherryKI/+^ embryos were dissected in PBS, and then dissociated in 0.05% trypsin (Wisent Cat. 325-042-CL) for 10 minutes at 37°C. Tryptic digest was terminated by adding 20% FBS, and cells were triturated ∼6 times before resuspending in PBS with 5 mM EDTA and 0.1% BSA. Dissociated cells (2.5 µl/1×10^6^ cells) were stained with Viability Dye (VD) eFluor 780 (eBioscience #65-0865-14) and with anti-CD15-Alexa Fluor 647 (BD Bioscience #560120) and CD15^+^, VD- and mCherry^+^ cells were sorted by Fluorescence-activated cell sorting (FACS). FACS sorted cells were then seeded at 1,000 cells/ml in DMEM/F12 (3:1), human FGF2 (40 ng/mL), human EGF (100ng/ml), B27 supplement (2%), N2 supplement (1%), Penicillin/streptomycin (0.1%), Fungizone (40 ng/mL), and 1 μM cyclopamine in 8 chamber slides coated with poly-d-lysine/laminin and cultured for 24 hrs. Chamber slides were prepared by coating with poly-D-lysine (10 ug/ml) in 1X PBS for 1 hr at room temperature, followed by coating with laminin (5 ug/ml in 1X PGS at 37°C for 3 hrs to overnight. Coated chamber slides were stored in PBS at 4°C until use. After 24 hrs, cells were fixed in 4% PFA in 1X PBS for 15 min at RT, washed 3 times with PBS and permeabilized with 0.2% triton X-100 for 10 min. Cells were blocked for 15 min at RT in blocking solution (10% horse serum in 1X PBS with 0.1% Triton X-100), and then incubated with primary antibodies diluted in blocking solution for overnight at RT. Antibodies included mouse anti-Tuj1 (β3-tubulin, 1:500, BioLegend #801202) and rabbit anti-Ki67 (1:500, Abcam #16667). Cells were then stained with secondary antibodies as described above, washed, and stained with DAPI before mounting in Aquapolymount. Two cell clones were analysed and considered to have undergone symmetric proliferative divisions if both daughters were Ki67^+^, symmetric differentiative divisions if both daughters were Tuj1^+^, and asymmetric divisions if there was a Ki67^+^ and Tuj1^+^ cell.

### Trajectory inference

scRNA-seq data of developing mouse neocortex was obtained from GSE153162 (Di Bella et al. 2021). Trajectory inference was performed on cells expressing Neurog1 and/or Neurog2 using the Monocle3 R package (Qiu et al. 2017). The input genes were top 4000 differentially expressed genes among Neurog1^+^, Neurog2^+^ and Neurog1-Neurog2 double^+^ cell populations using the differential GeneTest function with the adjusted p value cutoff < 0.001. The DDRTree algorithm was used for the inference.

### Gene regulatory network analysis (GRN) and in silico perturbation assay

We used a reconstructed GRN from our previous study (Fig. 5A of Han et al., 2021). Briefly, differentially expressed TFs (DETFs) were identified between the RNA-seq data of Neurog2 positive and negative NPC populations. Transcriptional interactions among these DETFs were obtained as the union of the E14.5 telencephalic Neurog2 ChIP-seq data (Sessa et al. 2017) and the MetaCore database (Nikolsky et al. 2005). Interactions were removed if the chromatin regions of target genes were not open in all ATAC-seq replicates. After building GRNs, in silico perturbation assays were performed using the program codes from (Okawa et al. 2015) by repressing the expression of either Neurog2 or Neurog1, or both together. GRN simulation was carried out by the Boolean network formalism using the inhibitor dominant logic rule. Boolean initial expression states 1 and 0 were assigned to differentially up- and down-regulated TFs, respectively. When GRN reached a steady state, repressed genes were identified. Cytoscape was used for visualization of GRNs (Shannon et al. 2003).

### Imaging, image analysis and statistics

P150 sections were imaged with a 10X objective (NA 0.45) for section overviews and a 20X objective (NA 0.75) for closer images, using the NikonA1 laser scanning confocal microscope system (Nikon Instruments, Melville, NY, USA). E12.5 and E15.5 images were acquired in one field of view (512 x 512 pixels), while E18.5 cortical columns were derived from tiled 512 x 512-pixel images. Postnatal images are presented as maximum intensity projections from Z-stacks since they are 30μm thick; embryonic images were acquired in one Z-plane only, as they were 10μm thick, which is the approximate size of a cell.

E12.5, 15.5, and 18.5 counting images were acquired using Leica Application Suite X (LAS X) on a Leica DMI8 fluorescent microscope from Leica Microscopy. All data are expressed as mean ± standard error of the mean (s.e.m.). One-way ANOVA with post hoc Tukey correction for multiple comparisons was used to compare cell numbers between more than two groups and an unpaired student’s t-test for comparisons of two groups. Statistical tests are indicated in the figure legends.

### Schematics

Schematics and Figures were generated using an Adobe Creative Cloud suite, including Illustrator V27.7 and Photoshop V24.6, or using an academic license to Biorender.

## ACKNOWLEDGEMENTS

This work was supported by an operating grant to CS from the Canadian Institutes of Health Research (CIHR) (PJT-162108). S.O. was supported by CORE grant from Fonds National de la Recherche Luxembourg (project number: R-AGR-3676-10-C). CS holds the Dixon Family Chair in Ophthalmology Research at the Sunnybrook Research Institute. AM was supported by scholarships from CIHR CGS-M, AMO was supported by scholarships from NSERC CGS-M, and SH from the Cumming School of Medicine, Ontario Graduate Scholarship (OGS), University of Toronto Vision Science Research Program, Peterborough K.M. HUNTER Charitable Foundation and Margaret and Howard GAMBLE Research Grant. We thank Justin You, Marielle Darnley (nee Balanaser) for technical assistance.

## AUTHOR CONTRIBUTIONS

AM, AMO, SO: conceptualization, formal analysis, investigation, methodology, visualization, validation, writing – original draft, writing – review and editing SH, LV, HG, DJD, DZ: formal analysis, investigation, writing – review and editing FG: resources, writing – review and editing. CS, AdS: funding acquisition, conceptualization, project administration, resources, supervision, validation, writing – original draft; writing – review and editing

